# Intraspecific trait variation shapes macro-scale patterns of niche breadth and range size

**DOI:** 10.64898/2026.10.31.763334

**Authors:** Ekadh Ranganathan, Philip D. Round, Juan C.T. Gonzalez, David V. Pluym, Luke L. Powell, Billi A. Krochuk, Chima J. Nwaogu, Charlotte M. Probst, Peter Mikula, Kwanye Z. Bitrus, Vedant Gattani, Andrew J. Pierce, Stephanie Y. Chia, James A. Villamin, Jarome R. Ali, Frane Babarović, Rolly C. Urizza, Kristof Zyskowski, Lukas J. Musher, Mohammad Irham, Brian C. Weeks, Sontaya Manawattana, Kyle D. Kittelberger, Charlene R. Bajamunde, Graciela D. Lorenzana, Rina R.B. Perez, Vanya G. Rohwer, Wangworn Sankamethawee, Camila G. Meneses, Robin Vijayan, Marianne L.K.Z. Fabella, Martin Bulla, Kier M. Pitogo, Woraphot Bunkhwamdi, Isabella C. Enriquez, Carla J. du Toit, Jestonie H. Namion, Sandupama Madhuhansi, Elizabeth Steell, Susana L. Mufumu, Nihan D. Dagtas, Amancio M. Etingue, Klara Widrig, Hari Basnet, Silvestre N.E. Malanza, Panagiotis Nikolaou, Anna M.B. Halili, Martin Hromada, Kelvin S.-H Peh, Maarten J.K. Hoepel, Caroline Dingle, Malcolm C.K. Soh, Daphawan Khamcha, Yue Wu, Michaël P.J. Nicolaï, René Marie Lafontaine, Olivier Pauwels, Florante A. Cruz, Peter A. Hosner, Alfredo Sánchez-Tójar, Joseph A. Tobias

## Abstract

Morphological variation within species may promote adaptation across ecological and geographic space, perhaps reflecting evolutionary plasticity and adaptability. Based on this concept, species with greater intraspecific variation in morphological traits may be expected to occupy broader ecological niches and larger geographic ranges. However, even in the age of global trait datasets, intraspecific morphological variation remains poorly quantified at broad taxonomic and geographic scales, limiting our understanding of how trait flexibility shapes ecological and biogeographic patterns. Focusing on 161 species of bulbuls, a widespread avian family containing some of the most abundant bird species in the world, along with some of the rarest and most restricted in range, we conducted global-scale sampling of morphological traits. These included beak shape, tarsus length and wing length, associated with dietary niche, foraging strategy and dispersal, respectively. Using phylogenetic models, we tested whether ecological niche breadth and geographical range size are predicted by intraspecific morphological variation quantified from 8,251 museum specimens and 5,627 live captures. Our results demonstrate that morphological trait variation predicts both ecological niche breadth and geographic range size. Furthermore, widespread species exhibit greater morphological variation than restricted-range species even when comparisons are constrained to equivalent geographic extents, consistent with the hypothesis that intraspecific trait variation facilitates range expansion rather than simply accumulating as a consequence of occupying a larger range. These results identify intraspecific morphological variation as a potentially important, yet largely overlooked, predictor of ecological traits and biogeographic distributions in birds.

## Introduction

Species differ markedly in their ecological niche sizes and geographic ranges, yet the factors that determine why some species occupy broad ranges while others remain geographically restricted and ecologically specialised are still poorly understood (Angert et al., 2020; Bolnick et al., 2003; Gaston, 2009; Radomski, 2026; Sexton et al., 2009). Morphological intraspecific trait variation (ITV), referring to differences in morphological traits among individuals within a species, has emerged as a potential mechanism underlying these patterns (Forsman and Wennersten, 2016; Sides et al., 2014; Valladares et al., 2014). By increasing the range of phenotypes expressed within a species, ITV may enhance ecological flexibility, enabling individuals to exploit a wider diversity of resources (Sides et al., 2014), tolerate broader environmental conditions (Valladares et al., 2014), and ultimately occupy broader niches and larger ranges (Bolnick et al., 2011; He et al., 2018).

Although morphological ITV has received increasing attention in ecology and evolution, research has focused predominantly on identifying the ecological and evolutionary drivers of ITV (Maldonado et al., 2017; Reinhold and Sánchez-Tójar, 2026; Van Valen, 1965) rather than testing whether ITV predicts macroecological outcomes (Pichancourt and Van Klinken, 2012). In particular, relatively little attention has been given to whether morphological ITV is associated with broader ecological niche breadth. Within birds, most studies have tested aspects of the Niche Variation Hypothesis (Van Valen, 1965), typically by relating morphological or dietary variation to niche breadth within populations or among only a small number of species (Johnson et al., 2022; Shaner et al., 2021). The only large comparative analysis spanning multiple avian families focused primarily on wing morphology, demonstrating that variation in wing length is associated with differences in feeding strategy (Reinhold and Sánchez-Tójar, 2026). Beyond birds, a growing body of evidence from plants indicates that populations exhibiting greater intraspecific functional trait variation tend to occupy broader environmental niches and tolerate a wider range of environmental conditions (He et al., 2018; Sides et al., 2014; Treurnicht et al., 2020). Together, these studies suggest that ITV may contribute to ecological flexibility across diverse taxa, but whether this relationship extends broadly across avian morphological traits remains unclear.

At the broader biogeographic scale, studies directly linking morphological ITV to species distributions also remain limited and have largely focused on variation in body mass or wing morphology (Fluck et al., 2026; Zulian and Youngflesh, 2025). Among the few studies addressing this question, Palacio et al. (2025) reported a positive association between beak length variation and the number of countries occupied by a species, while Delhey et al. (2013) found that species exhibiting greater plumage variation tended to have larger geographic distributions.

However, in a large-scale comparative study across birds, Zulian and Youngflesh (2025) found no significant association between morphological ITV and geographic range size. This apparent inconsistency likely reflects both the scarcity of comparative studies, and the traits examined. Morphological traits are unlikely to contribute equally to ecological flexibility, as wing morphology and body size may be more strongly constrained by aerodynamic and allometric selection, with wing morphology in particular showing evidence of evolutionary canalisation, whereby developmental and evolutionary constraints limit its capacity to vary in response to environmental conditions (Reinhold and Sánchez-Tójar, 2026). Due to this, any relationship between ITV and species distributions is likely to be trait dependent, yet comprehensive studies testing this pattern across diverse avian lineages remain largely absent.

An additional challenge is the inherent circularity between geographic range size and ITV. Species occupying larger ranges experience greater environmental heterogeneity, potentially generating elevated morphological variation through local adaptation or phenotypic plasticity (Chuang and Peterson, 2016; Cochrane et al., 2015; Yampolsky et al., 2014). As a result, even where positive associations are observed, it remains unclear whether increased ITV precedes range expansion or accumulates as species become more widespread.

Progress has also been limited by data availability (Moran et al., 2016). Reliable estimates of morphological ITV require dense sampling within species, yet most comparative studies in birds have relied on opportunistic bird-banding datasets (DeSante, 1992; Zulian and Youngflesh, 2025). Although invaluable for long-term ecological monitoring, these datasets were not designed for comparative analyses of ITV across entire clades, with species represented according to capture success rather than a targeted taxonomic sampling strategy, thereby limiting broad phylogenetic and macroecological inference. As such, ITV has rarely been quantified using a targeted deep-sampling framework spanning an entire avian clade, restricting our ability to investigate its predictive power at macroecological scales.

Bulbuls (*Pycnonotidae*) provide a powerful system for investigating the ecological outcomes of ITV, as the family comprises 161 largely non-migratory species spanning a remarkable spectrum of ecological strategies, from highly localised endemics with extremely restricted distributions to some of the most widespread and abundant passerines in the Old World (Fishpool and Tobias, 2005; Shakya and Sheldon, 2017). This diversity offers a rare opportunity to investigate why some closely related species appear inherently vulnerable to environmental change whereas others have successfully colonised a wide range of habitats across multiple continents. Here, we utilise this diversity by quantifying intraspecific morphological variation across the entire bulbul family using extensive museum and field specimen sampling. This study tests whether variation in multiple morphological traits predicts two fundamental macroecological properties, namely ecological niche breadth and geographic range size, and identifies which traits are most strongly associated with these outcomes (Fig. 1). Finally, to address the potential circularity between geographic range size and intraspecific variation, we compare widespread and restricted-range species variability within geographically matched subsets to determine whether elevated ITV persists in local populations, consistent with a role in promoting range expansion rather than just forming as a consequence of environmental heterogeneity.

**Figure 1:**
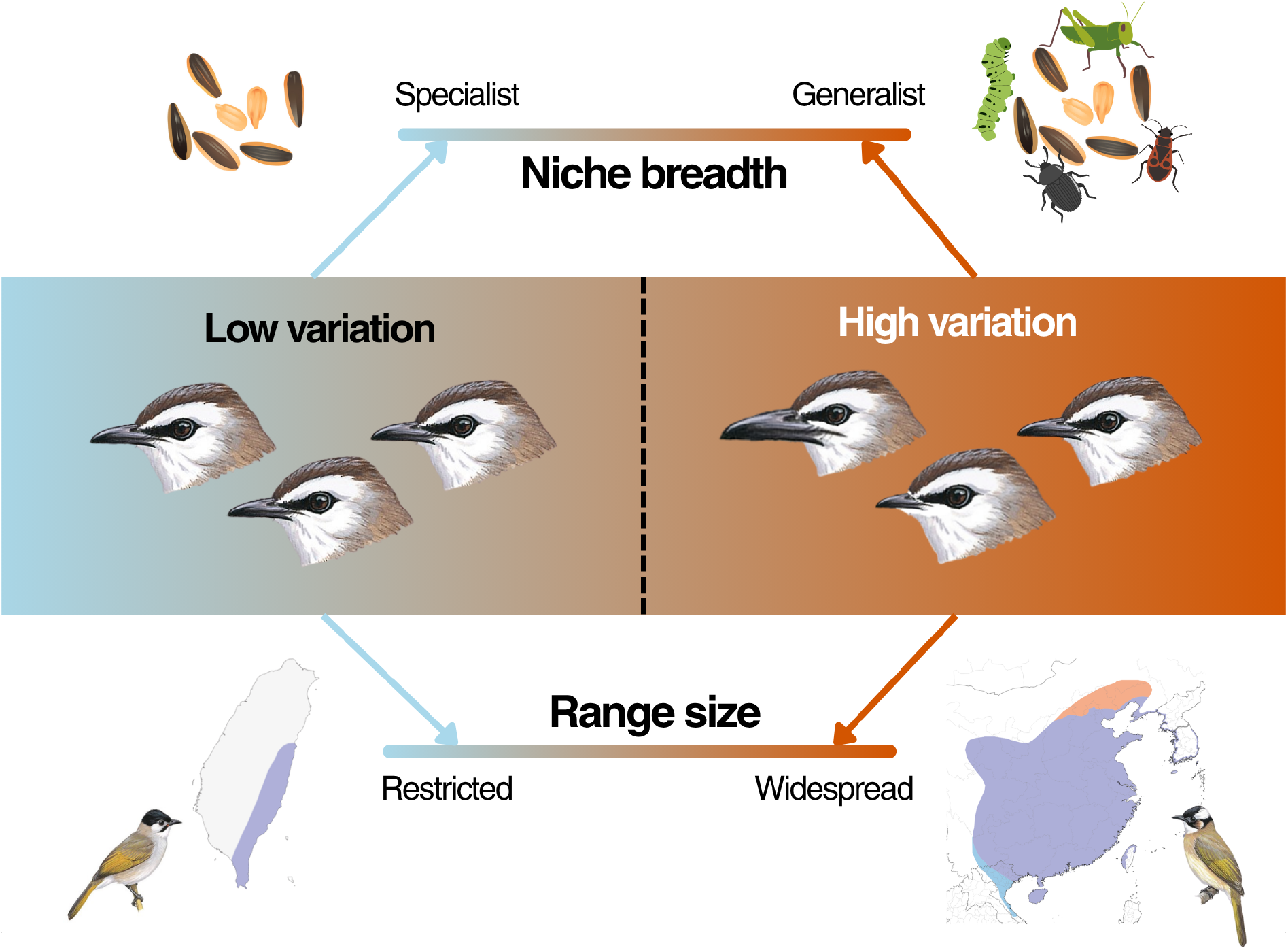
Conceptual figure summarizing the hypothetical link between high intraspecific variation and outcomes such as niche breadth and geographic range size. Under this view, variation between individuals reflects evolvability, or propensity to adapt to new or varied niches, which is linked to factors such as abundance and opportunistic foraging strategies, thereby permitting range expansion. This study tests the hypothesis that increasing intraspecific morphological variation promotes ecological flexibility, resulting in broader ecological niches and larger geographic ranges. The range maps at the bottom illustrate the contrast between a geographically restricted species (*Pycnonotus taivanus*; left) and its widespread close relative (*Pycnonotus sinensis*; right). Species illustrations have been reproduced with permission from the *Pycnonotus goiaiver*, *P.sinensis*, and *P.taivanus* accounts from *Birds of the World* (Cornell Lab of Ornithology).

## Results

Latent intraspecific morphological variation differed among traits. Mean species-level *CV_i_* values were highest for beak length (8.97%, ranging from 5.29% to 15.81%), followed by beak depth (6.74%, ranging from 5.03% to 9.15%), tarsus length (5.74%, ranging from 2.67% to 10.49%), and wing length (4.52%, ranging from 2.53% to 8.05%). Logged ecological niche breadth values ranged from 0.00 to 1.68 (mean = 0.75; Fig. 2, D), while geographic breeding range size varied by more than six orders of magnitude, ranging from 5.6 km^2^ to ∼ 19,600,000 km^2^, with a mean of ∼ 3,600,000 km^2^, indicating substantial variation across the family.

**Figure 2:**
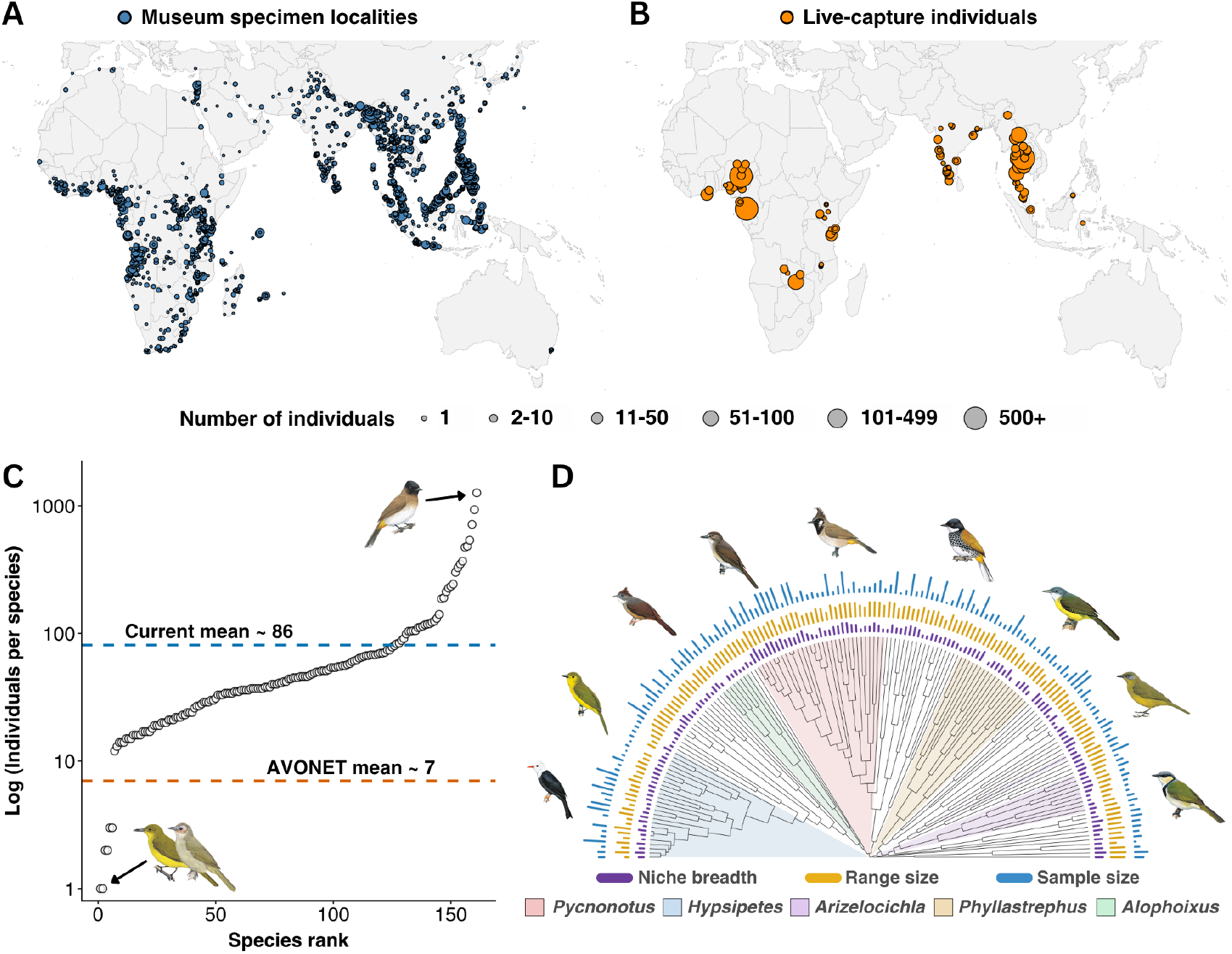
Broad geographic and phylogenetic sampling of bulbul morphology. (A) Geographic distribution of 2,684 unique specimen localities from museum collections, with point size proportional to the number of individuals sampled at each locality. (B) Geographic distribution of 153 unique localities at which live individuals were measured in the field, with point size proportional to the number of individuals sampled. (C) Number of individuals sampled per species, ranked from least to most sampled and shown on a logarithmic scale. The least-sampled species ware *Hypsipetes aureus* and *Nok hualon* (n = 1 individual), whereas the most-sampled species was *Pycnonotus barbatus* (n = 1262 individuals). Dashed lines indicate the mean number of individuals per species in AVONET (orange; ∼7 individuals; (Tobias et al., 2022)) and in the present dataset (blue; ∼86 individuals). (D) Phylogenetic distribution of the 161 bulbul species, based on the phylogeny of McTavish et al. (2025). Concentric bars show species-level niche breadth, geographic range size and specimen sampling effort. These variables were broadly distributed across the phylogeny rather than being concentrated within particular lineages. All bird illustrations were reproduced with permission from Cornell Lab of Ornithology through *Birds of the World*; scientific names for all illustrated species in panel D are provided in Supplementary Table 12, Supplementary Material.

Overall, latent intraspecific trait variation was positively associated with niche breadth and geographic range size, although the strength of these relationships differed among traits (Fig. 3). Greater variability in beak length (*β_CV_* = 0.42, 95% CrI = 0.15, 0.688), tarsus length (*β_CV_* = 0.281, 95% CrI = 0.07, 0.50), and beak depth (*β_CV_* = 0.78, 95% CrI = 0.36, 1.22) was associated with broader ecological niches, with posterior support of 99.9%, 99.5%, and 100.0%, respectively. Wing length variability showed little evidence of an association with niche breadth (*β_CV_* = 0.03, 95% CrI = −0.21, 0.27; posterior support = 59.5%).

**Figure 3:**
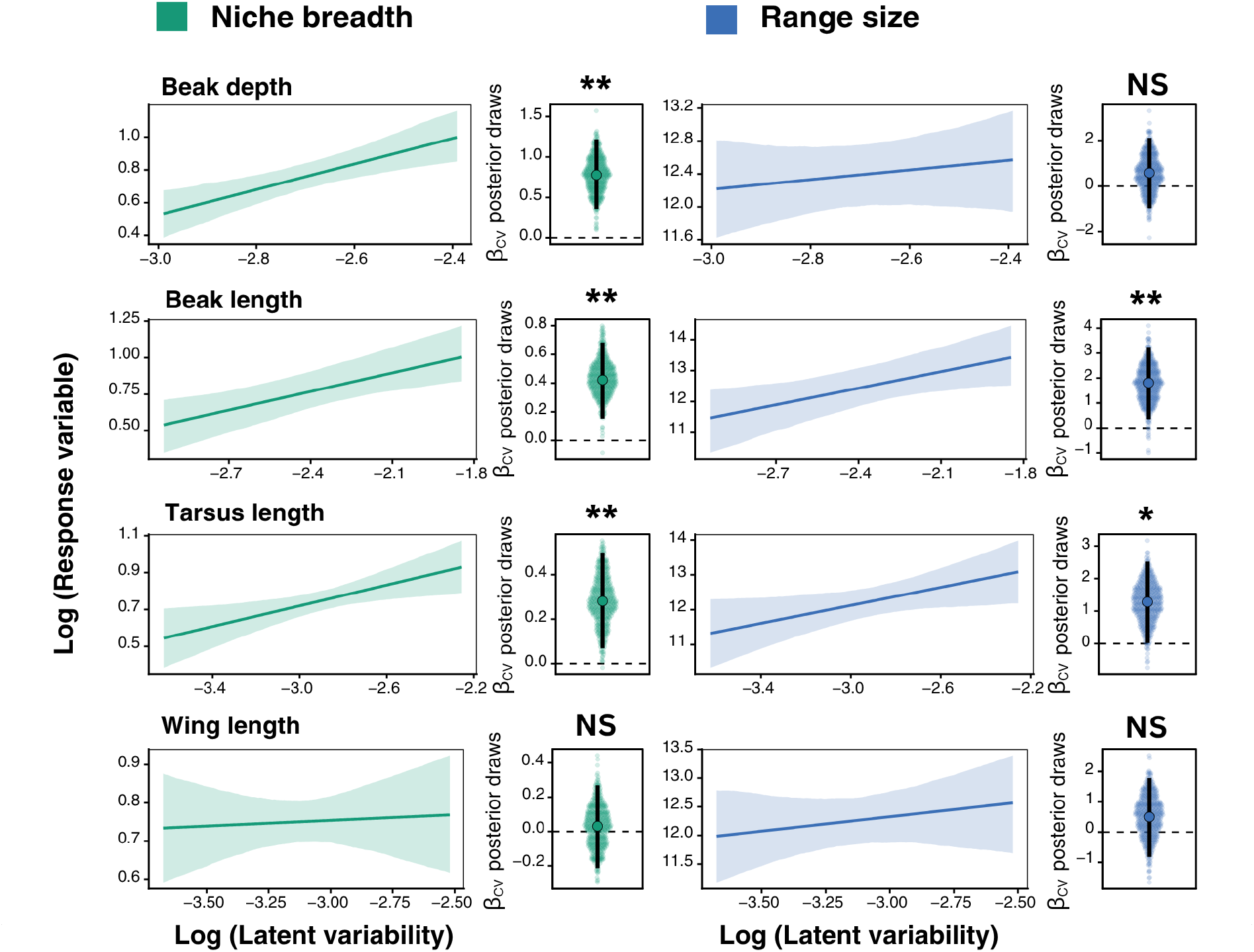
Intraspecific morphological variation as a determinant of niche breadth and geographic range size. Results shown are from phylogenetic Bayesian double hierarchical models (see Methods). ‘Beak length’ is measured tip-to-nares, which is more repeatable than culmen length across measurers (Tobias et al., 2022). For each morphological trait, the larger panels show the posterior mean relationship between latent trait variability and the response variable, with shaded ribbons indicating 95% credible intervals. Adjacent panels show the posterior distribution of the corresponding regression coefficient (*β_CV_*), with the posterior mean and 95% credible interval shown by the filled point and vertical error bar, respectively. Green panels represent models of niche breadth, whereas blue panels represent models of range size. Posterior support for the direction of each effect is indicated above the coefficient distributions, based on the proportion of posterior samples lying on one side of zero (**, ≥ 99% support; *, ≥ 95% support; NS (not significant), < 95% support).

Associations with geographic range size followed a similar pattern. Variability in beak length (*β_CV_* = 1.81, 95% CrI = 0.34, 3.24) and tarsus length (*β_CV_* = 1.28, 95% CrI = 0.02, 2.52) showed strong positive associations with geographic range size, with posterior support of 99.1% and 97.7%, respectively. In contrast, beak depth (*β_CV_* = 0.58, 95% CrI = −0.97, 2.11) and wing length (*β_CV_* = 0.50, 95% CrI = −0.82, 1.78) exhibited weaker positive relationships, with credible intervals overlapping zero and posterior support of 76.8% and 77.9%, respectively.

Because niche breadth and range size were weakly correlated, we repeated the analyses with latent trait variability modelled as the response variable and both ecological variables included simultaneously as predictors (Supplementary Table 2 in Supplementary Material). The positive associations between trait variability and geographic range size remained significant for beak length and tarsus length after accounting for ESI. Likewise, the associations between trait variability and ESI remained significant for beak length, beak depth and tarsus length after accounting for geographic range size as an additional fixed effect.

For the range-constrained paired-species analysis, the results presented use the finest spatial definition of a local population, corresponding to a 10 km clustering radius and a minimum of five specimens per locality cluster (Fig. 4). This dataset comprised 175 independent locality clusters for widespread species and 46 for their corresponding restricted-range relatives, with a mean of around ∼ 24 specimens per cluster.

**Figure 4:**
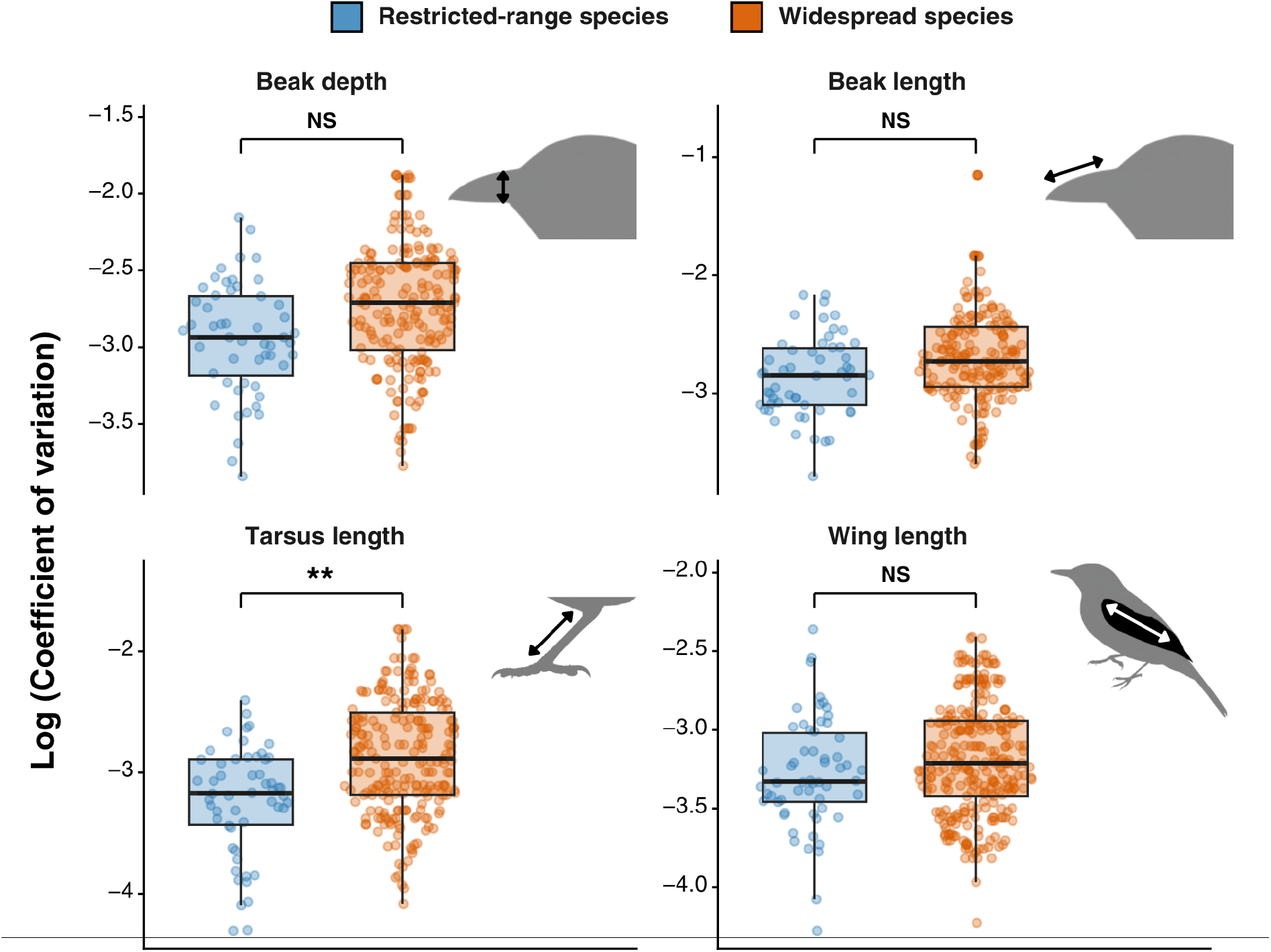
Range-constrained paired-species analysis comparing locality-level coefficients of variation for four morphological traits between geographically restricted and widespread bulbul species. Species were classified as widespread or restricted within clade-sister species pairs (Seddon et al., 2013) where the widespread species had a geographical range at least twice the size of its paired species. ‘Beak length’ is measured tip-to-nares, which is more repeatable than culmen length across measurers (Tobias et al., 2022). Individual points represent independent locality clusters identified using DBSCAN and are coloured by species range status (blue = restricted-range, orange = widespread). Boxplots show the median (central line), interquartile range (box), and non-outlier range (whiskers). Bi-directional arrows on the bird silhouettes indicate the dimension corresponding to the morphological measurement shown in each panel. Statistical significance of the range-status effect is indicated above each comparison (NS, not significant; **, *P* < 0.01). All bird illustrations were reproduced with permission from Cornell Lab of Ornithology through *Birds of the World*.

When comparing spatially matched local populations, widespread species consistently exhibited higher coefficients of variation than their restricted-range relatives, although the magnitude of this difference varied among traits. Locality-level variation in tarsus length (*β* = 0.257, *P* = 0.007) was significantly greater in widespread species, and although beak depth (*β* = 0.116, *P* = 0.127) and beak length (*β* = 0.094, *P* = 0.259) also showed positive effects for widespread range status, these relationships were not statistically supported. Similarly, although wing length variation was marginally greater in widespread species (*β* = 0.071), the effect was the weakest among all traits, and not statistically significant (*P* = 0.225).

## Discussion

This study demonstrates that morphological intraspecific variation (ITV) is associated with both ecological niche breadth and geographic range size in bulbuls, although the strength of these relationships differs among traits. Species exhibiting greater variability in traits associated with foraging morphology tended to occupy broader ecological niches and larger ranges. Furthermore, widespread species retained greater morphological variation than their restricted-range relatives at equivalent local spatial scales, suggesting that increased trait variation may not appear solely as a result of occupying broader and more environmentally heterogeneous ranges.

Our results demonstrate that the relationship between morphological ITV and ecological outcomes is strongly trait dependent. Variability in beak length, beak depth and tarsus length showed positive associations with ecological niche breadth, whereas wing length exhibited comparatively weak support.

One explanation is that the ecological consequences of morphological variation depend on the functional role of the trait. Traits that are evolutionarily labile and directly influence resource acquisition or habitat use may provide greater opportunities for populations to exploit a wider range of ecological conditions (Smith and Skúlason, 1996; Wennersten and Forsman, 2012). In bulbuls, variation in beak length, beak depth and tarsus length may reflect adaptation to an increased diversity of feeding strategies or foraging substrates available to individuals, thereby promoting the exploitation of a broader range of ecological conditions. In particular, the strong relationship between beak depth variability and niche breadth suggests that this trait may be more closely linked to ecological niche breadth than to geographic distribution, reflecting its importance in determining the diversity of food resources that individuals can exploit (Pigot et al., 2020; Reaney et al., 2020). These findings are consistent with theoretical predictions and empirical studies proposing that increased variation in traits associated with diet and habitat use should promote broader ecological niches through greater functional diversity within populations (Araújo et al., 2011; Pigot et al., 2020; Svanbäck and Schluter, 2012; Treurnicht et al., 2020).

Importantly, these trait-specific relationships were also evident when considering the broader biogeographic outcome of geographic range size. Variability in beak length and tarsus length showed the clearest positive associations with geographic range size, whereas beak depth and wing length exhibited comparatively weak support. The persistence of positive relationships for beak length and tarsus length across both niche breadth and geographic range size suggests that variation in these traits may contribute to ecological flexibility at multiple scales, from the utilisation of diverse resources and habitats to the occupation of geographically broader regions.

In contrast, wing morphology is likely to be subject to stronger stabilising selection because of its fundamental role in aerodynamic performance and flight efficiency, reducing the scope for within-species variation to accumulate (Baliga et al., 2019; Fitzpatrick, 1997; Reinhold and Sánchez-Tójar, 2026). The comparatively weak association between wing length variability and both ecological niche breadth and geographic range size may therefore reflect stronger functional constraints. Similarly, the weak relationship between beak depth variability and geographic range size, despite its strong association with niche breadth, suggests that the ecological effects of variation in this trait may be expressed more directly through resource use than through geographic occupancy. Thus, although morphological variability in most traits is associated with broader ecological niches, only some traits appear to translate this flexibility into the occupation of larger ranges.

The range-constrained paired-species analysis provides evidence that increased morphological ITV is not simply a consequence of occupying larger geographic ranges. If greater trait variation arose solely because widespread species experience a wider range of environmental conditions, then constraining comparisons to equivalent spatial scales would be expected to eliminate this relationship. Instead, widespread species consistently exhibited greater local-scale morphological variation than their restricted-range relatives, particularly for tarsus length. Although the strength of evidence varied among traits, the consistently positive direction of these differences suggests a broadly shared pattern rather than an effect confined to a single morphological trait. Furthermore, the trait-specific patterns recovered from the paired-species analysis closely followed those identified by the phylogenetic Bayesian model, with the strongest effects for tarsus length, followed by both beak traits, and the weakest for wing length. This consistency suggests that the same biological processes discussed above are likely to underlie patterns at the local scale as well.

Our findings are consistent with the possibility that increased ITV reflects an underlying capacity for evolutionary flexibility that may precede geographic range expansion. Previous studies have predominantly considered morphological variation as a consequence of environmental heterogeneity, asking how differences in climate or habitat generate ITV (Van Valen, 1965; Zulian and Youngflesh, 2025). In contrast, our paired-species analysis suggests that this variation is not just a consequence of widespread species experiencing greater environmental heterogeneity. Instead, elevated standing variation may reflect an inherent property of some species that provides greater scope for evolutionary responses to novel environmental conditions, potentially promoting colonisation of new regions and niche dimensions. This interpretation is broadly consistent with theoretical studies of range expansion, in which spatial sorting and natural selection can favour variation in ecologically important traits during colonisation of new habitats (Peischl and Gilbert, 2020; Phillips et al., 2010; Phillips and Perkins, 2019). Geographic expansion may nevertheless reinforce this process, as populations entering increasingly heterogeneous environments can experience further phenotypic diversification, generating a reciprocal eco-evolutionary feedback loop (Miller et al., 2020; Shine et al., 2011). Thus, increased morphological ITV may represent more than a by-product of occupying heterogeneous environments, as it may reflect an underlying capacity for evolutionary flexibility that predisposes some species to successful range expansion, with environmental heterogeneity subsequently reinforcing this variation.

Importantly, this conclusion proved robust to alternative definitions of a local population. The primary analyses, based on locality clusters with a 10 km radius and a minimum of five specimens, were quantitatively replicated even when the clustering radius was increased to 25 km (Supplementary Table 9 in Supplementary Material). Similar trends were also recovered using larger cluster radii and more stringent minimum sample sizes, although the association for tarsus length was weaker, while the association for beak length was statistically supported at a sampling radius of 50 km. However, these changes in support most likely reflect the substantial decrease in the number of locality clusters available for analysis under these stricter criteria, rather than a genuine weakening of the biological pattern. Collectively, these sensitivity analyses indicate that the inference of greater local-scale morphological variation in widespread species is not strongly dependent on the definition of a local population.

Despite these findings, the paired-species analysis should be viewed as an initial test of whether widespread species maintain consistently greater morphological ITV across local populations throughout their ranges, rather than as definitive evidence that elevated variation directly promotes range expansion. Although our sampling represents the most comprehensive specimen-level dataset currently available for an avian family, locality coverage remains opportunistic, with many species represented by relatively few well-sampled populations. Future studies combining denser geographic sampling across multiple populations and additional species-rich avian clades will be important for determining the generality of this pattern. In particular, explicitly comparing core and range-edge populations may provide further insight into how ITV is distributed across species ranges and whether elevated local variation truly characterises widespread species.

## Conclusions

This study provides strong evidence that intraspecific morphological variation is associated with both ecological niche breadth and geographic range size in a major avian family. Bulbul species exhibiting greater variation in traits associated with resource acquisition and foraging strategy consistently occupied broader ecological niches and larger geographic ranges, whereas variation in wing length showed little association with either macroecological property. Furthermore, morphological traits of widespread species retained greater variation than their restricted-range relatives at equivalent spatial scales, suggesting that elevated intraspecific variation is unlikely to evolve solely as a consequence of occupying larger and more heterogeneous ranges. The main alternative possibility is that high intraspecific variation reflects a form of underlying flexibility and evolvability, which may promote colonization of new regions and niche dimensions, thereby promoting range expansion. By combining deep specimen-level sampling across an entire avian family with hierarchical phylogenetic modelling, this study demonstrates that intraspecific morphological trait variation is a previously underappreciated predictor of macroecological patterns and highlights the importance of incorporating within-species variation into comparative studies of ecological and evolutionary diversification.

## Methods

### Data collection & validation

#### Assembling the specimen-level dataset

Intraspecific morphological variation was quantified from a global dataset of 13,878 specimens representing all 161 currently recognised bulbul species (Fig. 5, A). The dataset comprised 2,020 specimens obtained from the AVONET database (Tobias et al., 2022) and a further 11,858 specimens assembled specifically for this study. Of these additional specimens, 5,627 were measurements of living individuals, while 8,251 were specimens accessed by collaborators from 51 museum collections. Measurements across the complete dataset were contributed by 177 individual measurers.

**Figure 5:**
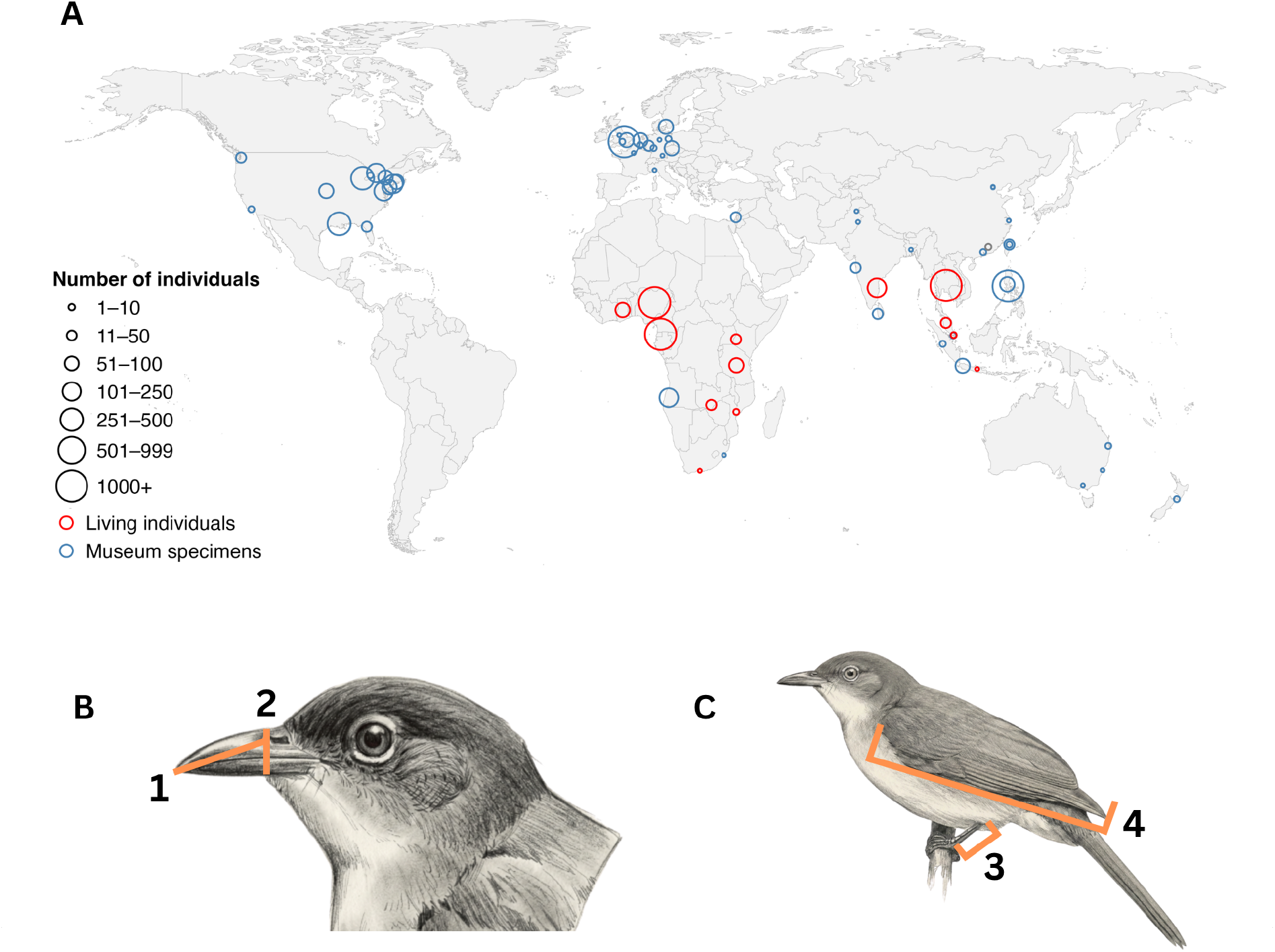
Overview of the specimen dataset and morphological measurements used in this study. (A) Global distribution of the 65 museum collections and field mist-netting projects contributing to the dataset. Circle size is proportional to the number of bulbul specimens measured at each location. Blue circles represent museum specimens, whereas red circles represent live birds measured in the field. (B) Beak measurements following the AVONET protocol: (1) beak length from the anterior edge of the nares to the bill tip, and (2) beak depth measured perpendicular to the anterior edge of the nares. (C) Additional morphological measurements: (3) tarsus length measured from the notch at the intertarsal joint (“knee”) to the third notch at the base of the toes (“ankle”), and (4) unflattened wing chord. Bird illustrations are reproduced with permission from the *Chlorocichla flaviventris* account from *Birds of the World* (Cornell Lab of Ornithology).

To standardise data collection across sources, all collaborators were provided with the AVONET measurement protocol (Tobias et al., 2022) and a standardised sampling sheet. All species names from resulting datasets were adjusted in line with AviList 2026 taxonomy (Rheindt et al., 2025) prior to assimilation into the main dataset to avoid naming incongruencies. Morphological measurements followed the AVONET protocol (Tobias et al., 2022), and included beak length from the anterior edge of the nares (tip-to-nares length), beak depth, tarsus length and unflattened wing chord length (Fig. 5, B–C). These traits were selected to maximise measurement repeatability while capturing complementary aspects of feeding, foraging and dispersal morphology. The tip-to-nares length (hereafter referred to as beak length) was the most repeatable measure of bill length among measurers (Tobias et al., 2022), beak depth was more repeatable than beak width, both of which are functionally associated with diet (Pigot et al., 2020), tarsus length provided the principal comparable measure of leg morphology in AVONET, and wing length is a highly repeatable measure of morphology associated with dispersal. Missing data percentages for each trait are reported in Supplementary Table 1 in Supplementary Material.

Quality control was performed separately for each species by flagging observations exceeding three standard deviations from the species mean and comparing them against the corresponding trait distributions in our dataset. Flagged records were inspected individually and excluded where measurement or transcription errors were identified.

#### Species-level niche breadth and geographic range estimates

Ecological niche breadth was quantified using the Ecological Specialisation Index (ESI) from BIRDBASE (Şekercioğlu et al., 2025), a species-level index derived as

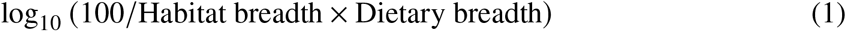

As lower ESI values indicate broader ecological niches, the variable was transformed to a positive measure of ‘niche breadth’ as 2 − ESI, such that higher values correspond to broader niches. Because habitat breadth is also strongly correlated with a species’ climatic and environmental niche (Barnagaud et al., 2012), this index captures an aspect of the Grinnellian niche through habitat use, and of the Eltonian niche through dietary breadth.

Species-level geographic range sizes were obtained from AVONET (Tobias et al., 2022), where breeding range area is reported in km^2^. Both geographic range size and niche breadth were log-transformed for analysis. Scientific names from both datasets were edited to align with AviList 2026 taxonomy, providing complete geographic range and niche breadth estimates for all currently recognised bulbul species.

### Modelling

#### Assembling the main model

Because the dataset consisted of both specimen-level morphometric measurements and species-level ecological predictors, we modelled the data using a phylogenetic Bayesian double hierarchical model (DHAM), built using the cmdstanr package (Gabry et al., 2025) in R Statistical Software (v4.5.1; R Core Team 2025). This framework jointly estimated species-specific trait variability from individual measurements while simultaneously modelling its relationship with species-level ecological variables. At the specimen level, the model accounted for measurement differences arising from data source (field versus museum specimens; Formula 2), whereas the species-level model estimated the association between latent trait variability and niche breadth or range size (Formula 3). This hierarchical structure allowed uncertainty arising at the specimen level to be propagated into estimates of species-level trait variability, and into the ecological relationships of interest.

Trait variability was represented by a latent species-specific coefficient of variation, denoted *CV_i_*. As with a conventional coefficient of variation, *CV_i_* is interpreted as the standard deviation relative to the species mean, representing the proportional deviation of trait values from the mean (i.e., *CV_i_* × 100 as a percentage). Expressing variation relative to the species mean also accounts for differences in body size among species, allowing the magnitude of trait variation to be compared independently of absolute trait size. However, unlike the conventional coefficient of variation calculated directly from species means and standard deviations, *CV_i_* was an estimated model parameter inferred from all specimen-level observations while accounting for measurement uncertainty and hierarchical structure. As such, species with fewer measurements were automatically shrunk towards the population mean, whereas estimates for well-sampled species were driven predominantly by their observed data.

For specimen *j* belonging to species *i*, trait values were modelled as

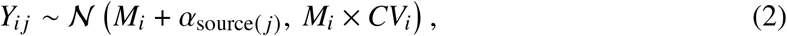

where *Y_i j_* is the observed trait value, *N* represents a normal distribution, *M_i_* is the estimated mean trait value for species *i*, *α*_source(_ *_j_*_)_ is the random effect of specimen data source (field or museum), and *M_i_* × *CV_i_* defines the species-specific standard deviation.

The latent trait variability *CV_i_* was then used as a predictor of each species-level response variable,

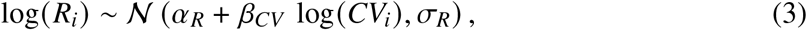

where *R_i_* denotes the species-level response (niche breadth or geographic range size), N represents a normal distribution, *α_R_* is the intercept, *β_CV_* is the main parameter of interest as it quantifies the association between latent trait variability and the response variable, while *σ_R_* represents the residual variation among species. *CV_i_* was log-transformed because its untransformed values were too small and departed from normality.

Because niche breadth and range size were expected to be correlated, we conducted an additional sensitivity analysis to determine whether their associations with latent trait variability were independent of one another. The main model was therefore repeated with latent trait variability modelled as the response variable and both log-transformed niche breadth and range size included simultaneously as fixed-effect predictors. This allowed the effect of each ecological variable to be evaluated while accounting for the other, thereby testing whether the observed relationships were robust to their underlying covariance. The results of this analysis are presented in Supplementary Table 2 in Supplementary Material.

Species-level trait means and latent coefficients of variation were modelled under a phylogenetic covariance structure using the recent phylogeny from McTavish et al. (2025), thereby accounting for shared evolutionary history between closely-related species.

Posterior inference was performed using Markov Chain Monte Carlo (MCMC) implemented in cmdstanr. Four independent Markov chains were run in parallel, each with 2,000 warm-up iterations followed by 4,000 sampling iterations. The target acceptance rate was set to 0.99 and the maximum tree depth to 15 to ensure sufficient exploration of the posterior distribution. Weakly informative priors were used throughout, comprising normal priors for intercepts and regression coefficients, standard normal priors for random effects, and exponential priors for variance parameters. Model convergence was assessed by confirming that all parameters had *R̂* < 1.01, and effective sample sizes were sufficient for reliable posterior estimation. An association was considered supported when the proportion of posterior samples lying on one side of zero (posterior support) was ≥ 95%.

#### Accounting for uneven sampling effort

Sample size differences across species were evaluated since specimen availability is inherently correlated with geographic range size, with widespread and abundant species generally being represented by larger museum collections than geographically restricted species. This made it difficult to disentangle whether increased trait variation was genuinely associated with geographic range size or simply reflected greater sampling effort. Because specimen sample size could not be included as a covariate in the main model without introducing substantial collinearity with geographic range size, we instead assessed whether reliable estimates of intraspecific variation could be recovered from relatively small numbers of specimens. To do this, we conducted a rarefaction analysis using species represented by more than 50 specimens (n = 70), in which the coefficient of variation for each trait was repeatedly estimated from nested random subsets of increasing sizes until estimates converged within ±1% of the value obtained from the full dataset. Across the four morphological traits, reliable estimates of the coefficient of variation were typically achieved with only 15–18 specimens per species (Supplementary Table 3 in Supplementary Material), indicating that estimates of trait variability stabilised beyond relatively modest sample sizes. As the majority of species included in the analyses exceeded these thresholds (n = 138 species exceeding 18 specimens), differences in specimen sampling depth are unlikely to have substantially influenced the estimated relationship between intraspecific variation and geographic range size.

#### Accounting for other potential biases

Although the main specimen-level model accounted only for data type (field versus museum specimens), other potential sources of measurement variation, including sex, measurer and locality, were evaluated separately through sensitivity analyses.

Sex was not included in the main model because approximately 45% of specimens lacked sex information, as sexual size dimorphism is generally weak across *Pycnonotidae* (Fishpool and Tobias, 2005). To assess whether sex influenced trait estimates, the main model was repeated using only specimens with known sex (M or F). The subsetted model showed that sex had a limited effect on trait measurements, as the *β_CV_* estimates were not significantly altered. The results of this sensitivity analysis are presented in Supplementary Table 4 in Supplementary Material.

Measurer was also excluded from the main model because sampling effort was highly uneven, with collaborators measuring a median of 12 specimens, at a median of only three species each. Including measurer as a species-specific random effect would therefore confound observer effects with species identity, particularly for narrowly distributed species represented by a single measurer. Instead, measurement consistency was quantified using a repeatability analysis based on the duplicate-specimen test described by Tobias et al. (2022), in which the same specimen was independently measured by different observers. A total of 62 pairs of duplicate measurements were included for this analysis, and repeatability coefficients were assessed using the rptR package (Stoffel et al., 2017). Across traits, all coefficients were > 0.9, indicating highly consistent measurements among measurers. The results of the repeatability analysis are presented in Supplementary Figure 2 and Supplementary Table 5 in Supplementary Material.

Locality was not included as a fixed or random effect in the main specimen-level model because geographic coordinates were unevenly distributed, with many collections concentrated at a small number of localities. Instead, we conducted a sensitivity analysis to test whether geographic separation among specimens influenced morphological differences within species. For each species and morphological trait, pairwise geographic distances between specimens were calculated and regressed against the corresponding absolute pairwise differences in trait measurements using ordinary least-squares regression. The resulting species-specific slopes quantified the relationship between geographic distance and morphological divergence, with slopes close to zero indicating little effect of geographic separation on trait variation. Across all traits, slope distributions were centred near zero and coefficients of determination were generally slightly positive but low, indicating that geographic distance explained little of the observed within-species morphological variation. The results are presented in Supplementary Figure 1 and Table **??** in Supplementary Material.

### Range-constrained paired-species test

To determine whether high intraspecific trait variation persists at local spatial scales in widespread species, we performed a range-constrained paired-species analysis. Following the approach of Seddon et al. (2013), 26 ‘clade-sister’ pairs were identified, with one species classified as widespread when its geographic range size was at least twice that of its geographically restricted relative (see Supplementary Table 7 in Supplementary Material for the selected species-pairs). Because some species formed sister relationships with more than one species in the phylogeny, individual species could contribute to multiple paired comparisons and were therefore not independent observations.

To compare morphological variation at equivalent spatial scales, specimens were georeferenced from locality strings using the GEOLocate software (Rios and Bart, 2010; see Supplementary Table 8 in Supplementary Material for more details on the georeferencing pipeline). Each species was then independently grouped into locality clusters using the density-based clustering algorithm DBSCAN implemented in the dbscan package (Hahsler et al., 2019). Cluster radii of 10, 25 and 50 km were evaluated to assess the sensitivity of the results to the definition of a local population, while a minimum cluster size of five specimens was chosen to ensure reliable estimates of intraspecific variation. For each retained cluster, coefficients of variation were calculated separately for each morphological trait.

Locality-level coefficients of variation were subsequently analysed using linear mixed-effects models fitted with restricted maximum likelihood using the lme4 package (Bates et al., 2015). For each trait, log-transformed coefficient of variation was modelled as a function of range status (widespread versus restricted), with log-transformed cluster sample size (*n*) included as a covariate to account for differences in sampling effort among local populations. At this spatial scale, coefficient of variation was used as a direct measure of local trait variability, as the smaller sample sizes within individual clusters meant that reliable estimation of a latent variability parameter like *CV_i_* would not be feasible. Random intercepts were included for both species-pair identity and species identity, yielding the model

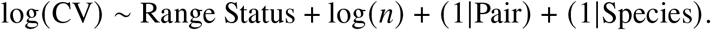

The ‘pair’ random effect allowed each comparison to have its own baseline level of morphological variation, whereas the species random effect accounted for repeated occurrences of the same species across multiple species-pair comparisons. Statistical significance of the range-status effect was assumed when *P* < 0.05.

## Data and Code Availability

Data and code generated for this study are available at

https://github.com/Ekadh/Bulbul-trait-variance.

Any additional information required to reanalyze the data reported in this paper is available from the main author upon request.

## Author Contributions

JAT and ER conceived the study. ER compiled the dataset, conducted the analyses, and wrote the manuscript. JC, PDR, DVP, LLP, BAK, CJN, CMP and PM also contributed substantial morphological datasets (> 500 specimens). JAT provided conceptual guidance and contributed to interpretation of the results.

## Acknowledgements

This study was made possible through the generous contributions of the Natural History Museum, Tring, UK, the University of the Philippines Los Baños, and 63 other museum collections and field projects worldwide, together representing 177 individual measurers who collected and measured the specimens used in this study. We thank all participating institutions, curators, collection managers, field researchers and volunteers for providing access to specimens and contributing morphometric measurements. A complete list of contributing collections and individual measurers is provided in Supplementary Table 10 and Supplementary Table 11 in Supplementary Material. We further want to thank Professor Alfredo Sánchez Tójar from the University of Coimbra for his invaluable insights during the data analysis phase of the project.

The authors also thank the Cornell Lab of Ornithology for permission to reproduce illustrations from *Birds of the World*, and all members of the Tobias Lab for helpful discussions throughout this project.

