## Supplementary Material for "Intraspecific trait variation shapes macro-scale patterns of niche breadth and range size"

This supplementary material contains additional methodological details, summary tables and sensitivity analyses results supporting the main text.

#### Supplementary Figures

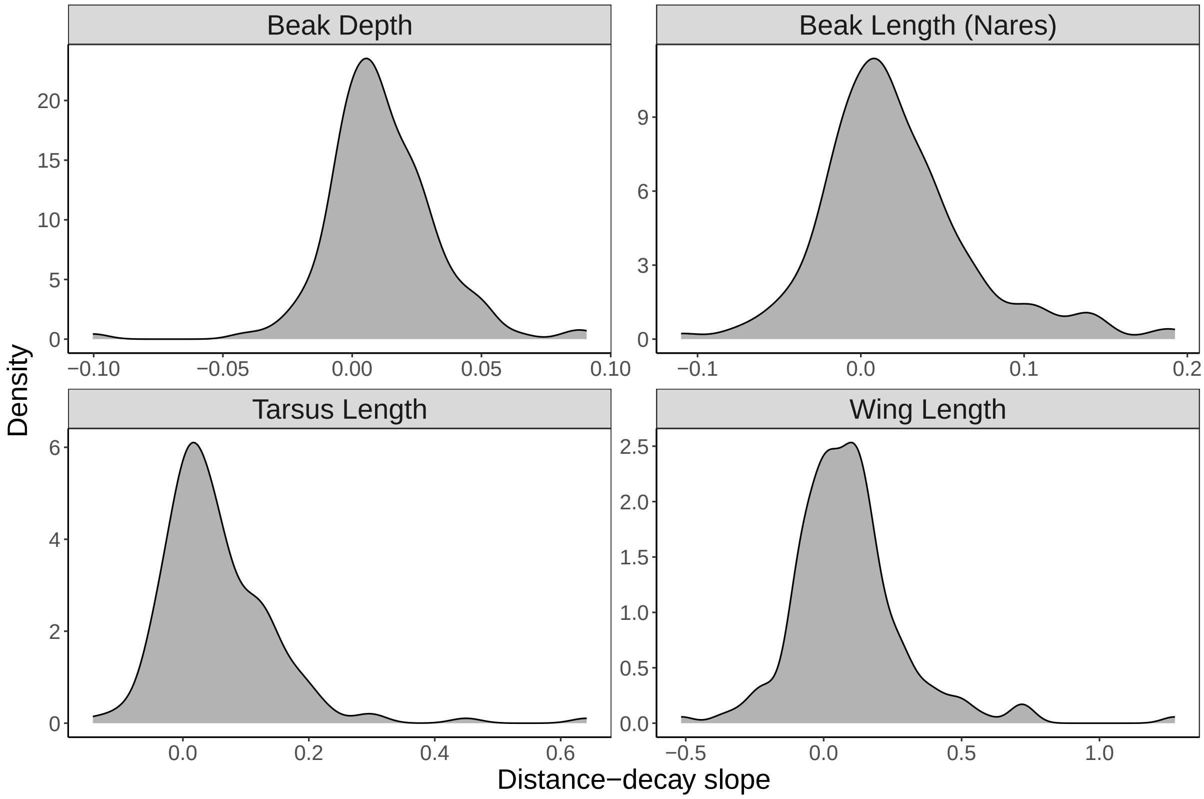


**Figure 1:** Species-specific distance-decay slope distribution plots for all bulbul species with > 2 unique sampling localities. Slope distributions are predominantly centered around zero with slight positive tails, indicating that the distance between sampling localities does not significantly affect trait measurements.

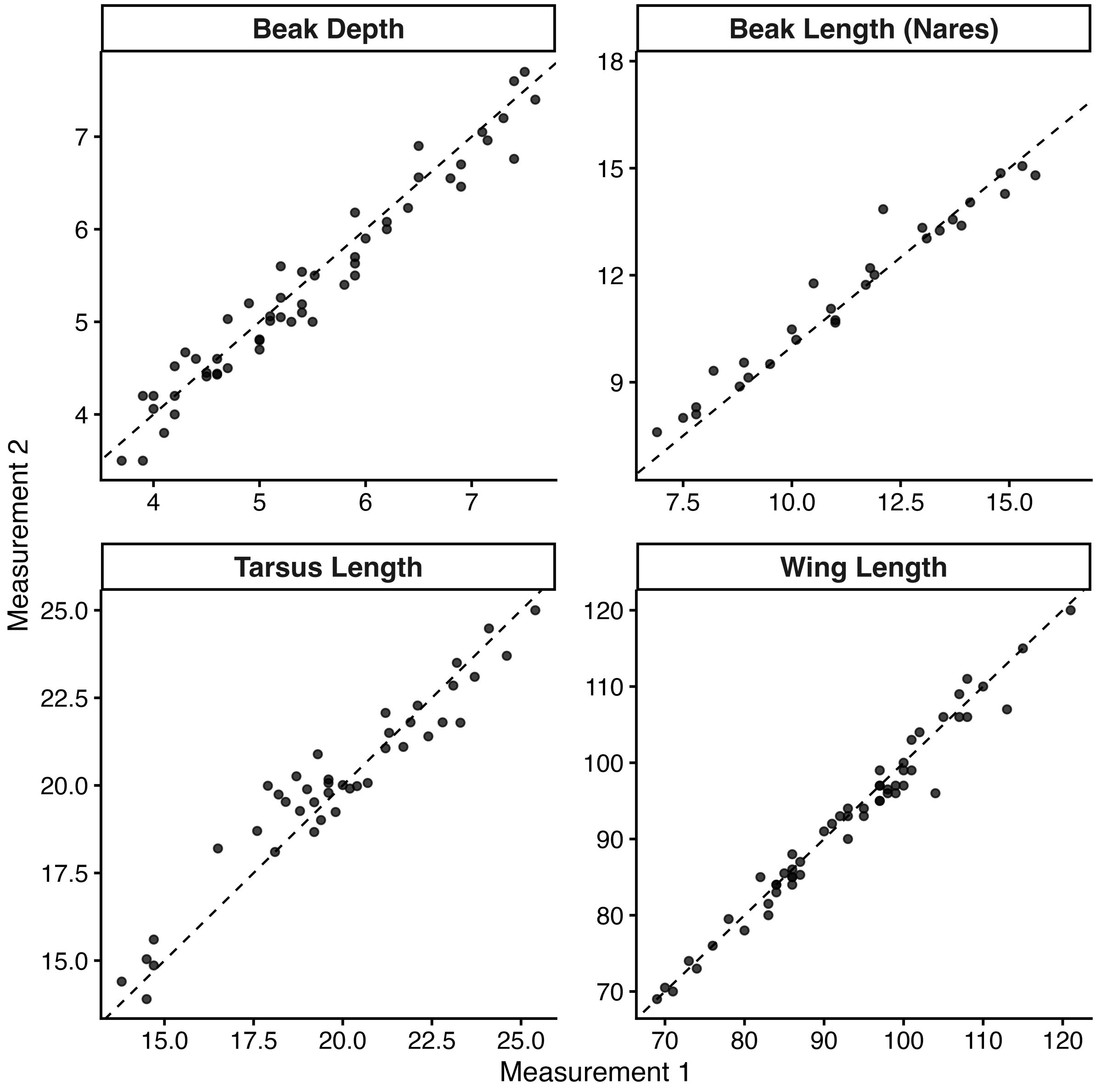


**Figure 2:** Agreement between duplicate measurements of the same specimens for the four focal morphological traits. Each point represents two independent measurements of a single specimen, and the dashed line indicates the 1:1 relationship expected under perfect agreement. The high concordance between repeated measurements demonstrates strong measurement consistency among observers across all traits.

### Supplementary Tables

**Table 1:** Missing data percentages for each morphological trait measured across all specimens.

| Morphological trait | Missing data (%) |
|---|---|
| Beak depth | 39.4 |
| Beak length | 40.9 |
| Tarsus length | 24.2 |
| Wing length | 2.6 |

**Table 2:** Results of the reverse Bayesian hierarchical model testing the independent effects of geographic range size and ecological specialisation on latent intraspecific trait variation after accounting for their covariance.

| Trait | Range size effect ($\beta$, 95% CrI) | ESI effect ($\beta$, 95% CrI) |
|---|---|---|
| Beak depth | 0.08 (0.05, 0.11) | 0.04 (0.00, 0.09) |
| Beak length | 0.07 (0.03, 0.10) | 0.06 (0.01, 0.11) |
| Tarsus length | 0.08 (0.04, 0.11) | 0.06 (0.00, 0.12) |
| Wing length | 0.08 (0.04, 0.11) | 0.00 (−0.05, 0.05) |

**Table 3:** Mean specimen sampling requirements estimated from the rarefaction analysis for each morphological trait. Values represent the mean number of specimens required for the estimated coefficient of variation to fall within ±1% of the true coefficient of variation across species.

| Trait | Mean specimens required | Species ($n$) | Tolerance to true CV |
|---|---|---|---|
| Beak depth | 16.67 | 70 | ±1% |
| Beak length | 16.09 | 70 | ±1% |
| Tarsus length | 17.83 | 70 | ±1% |
| Wing length | 15.14 | 70 | ±1% |

**Table 4:** Posterior estimates of the $\beta_{CV}$ slope parameter for the known-sex dataset, in which Sex was included as a fixed effect in the specimen-level model. Tarsus length was the only trait for which the 95% credible interval for the range size effect included zero in the known-sex model but not the main model. However, the posterior mean estimate (1.21) was comparable to that of the main model (1.28), suggesting that this difference in statistical support reflects increased uncertainty associated with the smaller known-sex dataset rather than a substantive change in the estimated effect.

| **Trait** | **Range size effect ($\beta$, 95% CrI)** | **ESI effect ($\beta$, 95% CrI)** |
|---|---|---|
| Beak depth | 0.59 (−1.00, 2.14) | 0.78 (0.34, 1.22) |
| Beak length | 2.36 (0.97, 3.71) | 0.505 (0.26, 0.76) |
| Tarsus length | 1.21 (−0.13, 2.48) | 0.239 (0.01, 0.48) |
| Wing length | 1.27 (−0.11, 2.64) | 0.24 (−0.04, 0.51) |

**Table 5:** Repeatability analysis based on 62 pairs of duplicate measurements of the same specimens by different observers. Repeatability ($R$) was estimated using a Gaussian mixed-effects model implemented in `rptR`, with specimen identity fitted as a random effect and 1,000 bootstrap replicates used to estimate 95% confidence intervals. Pearson's correlation coefficient ($r$) provides an additional measure of agreement between duplicate measurements.

| **Trait** | **Repeatability, $R$ (95% CI)** | **Pearson's $r$** |
|---|---|---|
| Beak depth | 0.971 (0.950, 0.982) | 0.975 |
| Beak length | 0.977 (0.958, 0.988) | 0.980 |
| Tarsus length | 0.947 (0.908, 0.970) | 0.960 |
| Wing length | 0.984 (0.975, 0.990) | 0.986 |

**Table 6:** Species-specific slope diagnostics for the relationship between logged pairwise geographic distance (km) and pairwise difference in trait measurements (mm).

| **Trait** | **Mean slope** | **Mean $R^2$** | **Mean $P$-value** | **Significant species** |
|---|---|---|---|---|
| Beak depth | 0.011 | 0.021 | 0.260 | 62 |
| Beak length | 0.022 | 0.017 | 0.281 | 57 |
| Tarsus length | 0.056 | 0.028 | 0.214 | 87 |
| Wing length | 0.092 | 0.015 | 0.262 | 62 |

**Table 7:** Complete list of clade-sister species pairs included in the range-constrained paired-species analysis. Widespread species were defined as having a geographic range at least twice that of their paired restricted species.

| Pair | Restricted species | Widespread species |
|---|---|---|
| 1 | *Alophoixus flaveolus* | *Alophoixus pallidus* |
| 2 | *Alophoixus frater* | *Alophoixus tephrogenys* |
| 3 | *Arizelocichla tephrolaema* | *Arizelocichla masukuensis* |
| 4 | *Atimastillas flavicollis* | *Atimastillas flavigula* |
| 5 | *Bleda notatus* | *Bleda canicapillus* |
| 6 | *Chlorocichla falkensteini* | *Chlorocichla flaviventris* |
| 7 | *Criniger olivaceus* | *Criniger calurus* |
| 8 | *Hemixos cinereus* | *Hemixos flavala* |
| 9 | *Hypsipetes mindorensis* | *Hypsipetes philippinus* |
| 10 | *Hypsipetes olivaceus* | *Hypsipetes madagascariensis* |
| 11 | *Hypsipetes siquijorensis* | *Hypsipetes philippinus* |
| 12 | *Hypsipetes thompsoni* | *Hypsipetes leucocephalus* |
| 13 | *Microtarsus fuscoflavescens* | *Microtarsus melanocephalos* |
| 14 | *Phyllastrephus albigula* | *Phyllastrephus debilis* |
| 15 | *Phyllastrephus fischeri* | *Phyllastrephus cabanisi* |
| 16 | *Phyllastrephus poliocephalus* | *Phyllastrephus albigularis* |
| 17 | *Phyllastrephus viridiceps* | *Phyllastrephus albigularis* |
| 18 | *Pycnonotus capensis* | *Pycnonotus barbatus* |
| 19 | *Pycnonotus davisoni* | *Pycnonotus finlaysoni* |
| 20 | *Pycnonotus leucogenys* | *Pycnonotus leucotis* |
| 21 | *Pycnonotus nigricans* | *Pycnonotus barbatus* |
| 22 | *Pycnonotus snouckaerti* | *Pycnonotus bimaculatus* |
| 23 | *Pycnonotus taivanus* | *Pycnonotus sinensis* |
| 24 | *Rubigula gularis* | *Rubigula flaviventris* |
| 25 | *Rubigula montis* | *Rubigula dispar* |
| 26 | *Rubigula squamata* | *Rubigula erythropthalmos* |

**Table 8:** Classification scheme used to georeference specimen localities based on the information available in locality descriptions.

| Locality string type | Example locality string | Action taken when assigning coordinates |
|---|---|---|
| Exact locality / offset | "5 km N of X", GPS coordinates, named site | Placed coordinate at the exact locality. |
| Small village / town | "Libokwa village", "Etawah town" | Placed coordinate at the geographic centre of the locality. |
| City / National Park / Small Island | "Bangalore", "Accra", "Serengeti National Park", "Sabtang Island" | Placed coordinate at the geographic centre of the locality. |
| District / County / River | "Coimbatore District", "River Dja" | Placed coordinate at the geographic centre of the locality. |
| State / Region | "Tamil Nadu", "Northern Province" | Placed coordinate at the species' range centroid within the region. |
| Large island / Country | "Borneo", "Madagascar", "Java" | Placed coordinate at the species' range centroid within the region. |

**Table 9:** Sensitivity analysis of the range-constrained paired-species model using a 25 km locality clustering radius. Estimates represent the effect of widespread species relative to their geographically restricted sister species on logged coefficients of variation after accounting for sample size. Bolded values indicate statistically significant results.

| Trait | Estimate | SE | *P* |
|---|---|---|---|
| Beak depth | 0.095 | 0.074 | 0.205 |
| Beak length | 0.089 | 0.082 | 0.283 |
| Tarsus length | 0.221 | 0.094 | **0.022** |
| Wing length | 0.026 | 0.054 | 0.635 |

**Table 10:** The data sources contributing specimens to this study, comprising 51 museum collections and two types of field-based datasets. Individual field and museum specimen measurers are acknowledged separately in Supplementary Table 11.

| Source | Institution | Specimens | City | Country |
|---|---|---|---|---|
| NHMUK | Natural History Museum | 1576 | London | UK |
| UPLB | University of the Philippines Los Baños, Museum of Natural History | 1047 | Los Baños | Philippines |
| UMMZ | University of Michigan, Museum of Vertebrate Zoology | 688 | Ann Arbor | USA |
| LSUMZ | Louisiana State University, Museum of Natural Science | 584 | Baton Rouge | USA |
| ISCED | Instituto Superior de Ciências de Educação da Huíla (ISCED-Huíla) | 488 | Lubango | Angola |
| ROM | Royal Ontario Museum, Department of Natural History | 452 | Ontario | Canada |
| USNM | Smithsonian Institute National Museum of Natural History | 360 | Washington DC | USA |
| AMNH | American Museum of Natural History | 301 | New York | USA |
| PNM | Philippine National Museum | 234 | Manila | Philippines |
| KU | University of Kansas Biodiversity Institute | 228 | Lawrence | USA |
| NHMD | Natural History Museum of Denmark, University of Copenhagen | 218 | Copenhagen | Denmark |
| YPM | Yale Peabody Museum | 217 | New Haven | USA |
| RMNH | Naturalis Biodiversity Center | 206 | Leiden | Netherlands |
| MZB | Museum Zoologicum Bogoriense, National Research and Innovation Agency (BRIN) | 204 | Bogor | Indonesia |
| ANSP | Academy of Natural Sciences of Drexel University (Philadelphia) | 183 | Philadelphia | USA |
| UMZC | University Museum of Zoology, Cambridge | 173 | Cambridge | USA |
| CUMV | Cornell University Museum of Vertebrates | 122 | Ithaca | USA |
| NMP | National Museum in Prague | 113 | Prague | Czechia |
| NMC | National Museum of Colombo | 86 | Colombo | Sri Lanka |
| SMNHTAU | The Steinhardt Museum of Natural History | 83 | Tel Aviv | Israel |

*Table 10 continued*

| Source | Institution | Specimens | City | Country |
|---|---|---|---|---|
| ZFMK | Zoologisches Forschungsmuseum Alexander Koenig | 79 | Bonn | Germany |
| BNHS | Bombay Natural History Society | 70 | Mumbai | India |
| FLMNH | Florida Museum of Natural History | 65 | Gainesville | USA |
| TBRI | Taiwan Biodiversity Research Institute | 56 | Jiji | Taiwan |
| UWBM | University of Washington, Burke Museum of Natural History and Culture | 54 | Seattle | USA |
| SYS | Museum of Biology, Sun Yat-sen University | 42 | Guangzhou | China |
| OUM | Oxford University Museum of Natural History | 41 | Oxford | UK |
| RBINS | Royal Belgian Institute of Natural Sciences | 31 | Brussels | Belgium |
| WFVZ | Western Foundation of Vertebrate Zoology | 29 | Camarillo | USA |
| QM | Queensland Museum | 25 | Brisbane | Australia |
| UWO | University of Western Ontario | 23 | London | Canada |
| LKCNHM | Lee Kong Chian Natural History Museum, National University of Singapore | 23 | Singapore | Singapore |
| MZUA | Museum Zoologi Universitas Andalas | 19 | Padang | Indonesia |
| SMF | Senckenberg Forschungsinstitut und Naturmuseum | 17 | Frankfurt | Germany |
| BRCAS | Biodiversity Research Museum, Academia Sinica, Taiwan | 16 | Taipei | Taiwan |
| NMNZ | Museum of New Zealand Te Papa Tongarewa | 16 | Wellington | New Zealand |
| ZMB | Leibniz-Institut für Evolutions und Biodiversitätsforschung an der Humboldt-Universität zu Berlin | 11 | Berlin | Germany |
| IZCAS | Institute of Zoology, Chinese Academy of Sciences | 10 | Beijing | China |
| AM | Australian Museum | 10 | Sydney | Australia |
| NHM-BS | Staatliches Naturhistorisches Museum | 6 | Braunschweig | Germany |
| UO | University of Okara | 6 | Okara | Pakistan |
| KZHM | Emeritus Professor Kazi Zaker Husain Museum | 5 | Dhaka | Bangladesh |
| NMV | Museums Victoria | 5 | Melbourne | Australia |
| MNHN | Muséum national d’Histoire naturelle | 4 | Paris | France |

*Table 10 continued*

| Source | Institution | Specimens | City | Country |
|---|---|---|---|---|
| PMNH | Pakistan Museum of Natural History | 4 | Islamabad | Pakistan |
| MMUM | Manchester Metropolitan University Museum | 3 | Manchester | UK |
| DNSM | Durban Natural Science Museum | 2 | Durban | South Africa |
| SNHM | Shanghai Natural History Museum | 2 | Shanghai | China |
| SNSB-ZSM | Staatliche Naturwissenschaftliche Sammlung Bayerns – Zoologische Staatssammlung München | 2 | München | Germany |
| BMNHC | Beijing Museum of Natural History | 1 | Beijing | China |
| MSNG | Natural History Museum of Giacomo Doria | 1 | Genova | Italy |
| Field 1 | Unpublished data from multiple field projects | 5627 | – | – |
| Field 2 | Published live specimen dataset (?) | 10 | – | – |
| **Total** | | **13,878** | | |

**Table 11:** Individual measurers, their corresponding identifiers, and the number of specimens measured by each observer. These records include both individual and collaborative measurement entries.

| Measurer | Measurer name | Individuals measured |
|---|---|---|
| AAB | Adham Ashton-Butt | 2 |
| AD | Adam J.M. Devenish | 18 |
| AJP | Andrew Pierce | 339 |
| AK | Adi Karya | 5 |
| AKL | Aung K. Lin | 1 |
| AKS | Akshay Herur | 9 |
| AKW | Andrew Wiegardt | 94 |
| AL | Alan Leishman | 8 |
| ALN | Albert Noorlander | 1 |
| AME | Amancio M. Etingue | 80 |
| AMH | Anna M.B. Halili | 51 |

*Table 11 continued*

| Measurer | Measurer name | Individuals measured |
|---|---|---|
| APA | Aparna Rao | 1 |
| ARP | Arpitha Bhat | 8 |
| ARS | Archita Sharma | 25 |
| AW | Ashwin Warudkar | 24 |
| BAK | Billi Krochuk | 70 |
| BJJ | Bashier Jr. Jailani | 19 |
| BP | Ben Phalan | 2 |
| BRW | Brian C. Weeks | 157 |
| CA | Chiti Arvind | 6 |
| CAM | Camila Meneses | 118 |
| CC | Chanon Chirachitmichai | 1 |
| CDT | Carla du Toit | 89 |
| CJD | Caroline Dingle | 54 |
| CJN | Chima Nwaogu | 536 |
| CMP | Charlotte M. Probst | 531 |
| CN | Chukiat Nualsri | 7 |
| CRB | Charlene R. Bajamunde | 97 |
| CRB/MLF | Charlene R. Bajamunde/Marianne L.K.Z. Fabella | 42 |
| CSM | Coloma S.N. Malabo | 3 |
| DA | Desmond Allen | 2 |
| DK | Daphawan Khamcha | 52 |
| DKO | Daniel Kong | 5 |
| DML | Dayna Levine | 4 |
| DNA | Desmond Allen | 46 |
| DVP | David V. Pluym | 584 |
| EH | Enam Ul Haque | 7 |

*Table 11 continued*

| Measurer | Measurer name | Individuals measured |
|---|---|---|
| EMS | Elizabeth Steell | 84 |
| ER | Ekadh Ranganathan | 997 |
| FZ | Faizee A. Khan | 1 |
| GB | Gerard Brett | 1 |
| GBE | Graham Beal | 1 |
| GDL | Graciela D. Lorenzana | 137 |
| GDR | Glaucia Del-Rio | 32 |
| GG | George Gale | 15 |
| GJI | Greg J. Irving | 3 |
| GPM | Geoff Mawson | 5 |
| GTI | Gayathri R. | 1 |
| HB | Hannah Batten | 28 |
| HBA | Hari Basnet | 75 |
| HD | Henrik Dahl | 1 |
| ICE | I.C. Enriquez | 103 |
| IJ | Israt Jahan | 5 |
| JALV | James A.L. Villamin | 335 |
| JAT | Joseph A. Tobias | 36 |
| JC | Juan C.T. Gonzalez | 110 |
| JDT | Jaden Tran | 23 |
| JEM | Jonathan Murray | 32 |
| JHN | Jestonie H. Namion | 84 |
| JK | Jitwadee Khoonwongsa | 41 |
| JND | Nick Dymond | 9 |
| JRA/FB | Jarome R. Ali/Frane Babarovic | 266 |
| JS | James Steward | 5 |

*Table 11 continued*

| Measurer | Measurer name | Individuals measured |
|---|---|---|
| JV | Jan Visser | 9 |
| KAP | Kacharat Phromkunathon | 2 |
| KBZ | Kwanye Z. Bitrus | 478 |
| KEK | Kevin Kelly | 1 |
| KHT | Khemthong Tonsakulrungruang | 10 |
| KI | Kultida Itthiporn | 12 |
| KJT | Kevin Thornton | 8 |
| KKB | Kyle D. Kittelberger | 141 |
| KMP | Kier M. Pitogo | 110 |
| KP | Korakoch Pobprasert | 31 |
| KS | Kaset Sutasha | 1 |
| KSH | Kaushlendra Singh | 14 |
| KT | Kihoko Tokue | 44 |
| KW | Klara Widrig | 77 |
| KZ | Kristof Zyskowski | 217 |
| KZT | Kyaw Z. Tun | 2 |
| LAB | Lainie Berry | 5 |
| LB | Laura Kettering (née Birkmann) | 23 |
| LJM | Lukas J. Musher | 183 |
| LLP | Luke L. Powell | 60 |
| LLP/BAK | Luke L. Powell/ Billi Krochuk | 554 |
| MB | Martin Bulla | 113 |
| MBN | Mohammed Mubeen | 1 |
| MCKS | Malcolm C.K. Soh | 53 |
| MFJ | Mike Jackson | 1 |
| MFZ | Michael A. Ford | 16 |

*Table 11 continued*

| Measurer | Measurer name | Individuals measured |
|---|---|---|
| MH | Maarten Hoepel | 56 |
| MHY | Muhammad Tayyub | 10 |
| MLF | Marianne L.K.Z. Fabella | 35 |
| MMU | Martha Mutiso | 3 |
| MNE | Maria N.E. Nchama | 11 |
| MNJ | Muhammad N. Janra | 19 |
| MOI | Mohammad Irham | 192 |
| MOS | Moses Chibesa | 7 |
| MP | Martin Päckert | 8 |
| MPJN | Michaël P.J. Nicolaï | 49 |
| MPM/MLF | Ma. Patricia M. Moralde/Marianne L.K.Z. Fabella | 18 |
| MRH | Martin Hromada | 64 |
| MSU | Maha S. Unii | 7 |
| MXK | Max Khoo | 1 |
| ND | Nick Dymond | 2 |
| NDD | Nihan D. Dagtas | 83 |
| NG | Naman Goyal | 36 |
| NGL | Ngwe Lwin | 3 |
| NJC | Nigel Collar | 227 |
| NWR | Nick W. Ramsey | 1 |
| OAC | Orlando Acevedo-Charry | 33 |
| OAJ | Omarkayam A. Jaafar | 42 |
| PCG | Patricia C. Guedes | 4 |
| PD | Poramad Dougkomnna | 3 |
| PDR | Philip Round | 337 |
| PET | Phetprakai | 22 |

*Table 11 continued*

| Measurer | Measurer name | Individuals measured |
|---|---|---|
| PFR | Patricia F. Rodrigues | 47 |
| PLK | Paisin Lekcharoen | 23 |
| PM | Peter Mikula | 513 |
| PMU | Paul A.M. Umale | 21 |
| PN | Panagiotis Nikolaou | 74 |
| POL | Tun Tun Oo | 4 |
| PW | Patrick A. Walkden | 38 |
| RA | Rongrong Angkaew | 5 |
| RAB | Rob Barber | 5 |
| RB | Raja S. Bandi | 8 |
| RBP | Rina R.B. Perez | 127 |
| RC | René Corado | 1 |
| RIF | Richard Fuller | 25 |
| RK | Rungsrit Kanjanavanit | 1 |
| RL | Richard Littauer | 16 |
| RML/OP | René Marie Lafontaine/Olivier Pauwels | 31 |
| ROC | Ross Crates | 2 |
| RV | Robin Vijayan | 116 |
| SAI | Chadaporn Srisai | 1 |
| SAL | Susana Lin | 84 |
| SAM | Sandupama Madhuhansi | 86 |
| SEM | Silvestre N.E. Malanza | 75 |
| SGS | Stephen Samworth | 14 |
| SHA/MMR | Shofiul Alam/Md. Mizanur Rahman | 5 |
| SHM | Saw H. Mim | 1 |
| SHV | Shivam | 17 |

*Table 11 continued*

| Measurer | Measurer name | Individuals measured |
|---|---|---|
| SK | Somboon Kamtaeja | 5 |
| SN | Somchai Nimnuan | 36 |
| SON | Sontaya Manawattana | 155 |
| ST | Supatcharee Tanasarnpaiboon | 19 |
| SUB | Subash | 1 |
| SUP | Supat Prasobsin | 6 |
| SYC | Stephanie Chia | 339 |
| TB | Tom Backlund | 2 |
| TE | Ton Eggenhuizen | 3 |
| TEM | Pornkasem Kantamara | 1 |
| THU | Thura S.M. Htike | 1 |
| TO | Tiwa Ong-in | 4 |
| TP | Thanathip Panyawutthitham | 1 |
| TSK | Thanyarat Sukruan | 2 |
| VG | Vedant Gattani | 452 |
| VGR | Vanya G. Rohwer | 122 |
| VIJ | Viral Joshi | 19 |
| VK | Varun Kher | 15 |
| VKL | Vinay K.L. | 8 |
| VNJA | Venus N.J. Abacahin | 16 |
| VNJA/MLF | Venus N.J. Abacahin / Marianne L.K.Z. Fabella | 21 |
| VNU | Vishnudas C.K. | 42 |
| VS | Vattikorn Sophonrat | 4 |
| VWA | Vishwa Jagati | 4 |
| WAS | Wasana | 2 |
| WB | Woraphot Bunkhwamdi | 107 |

*Table 11 continued*

| Measurer | Measurer name | Individuals measured |
|---|---|---|
| WCH | Warut Chaleekarn | 2 |
| WJ | Bill Jones | 8 |
| WL | Wichyanan Limparungpatthanakij | 17 |
| WS | Wangworn Sankamethawee | 120 |
| WZ | Wenyi Zhou | 32 |
| X | Mikio Nayohara | 1 |
| YD | Yash Deshpande | 3 |
| YUP | Yupin | 1 |
| YW | Yue Wu | 52 |
| YYT | Yen Yi Tan | 37 |
| ZWE | Zwe Zaw Zaw Hein | 1 |

**Table 12:** Scientific names of species included in Fig. **??**, from left to right, reproduced with permission from *Birds of the World*, Cornell Lab of Ornithology.

| Order | Species |
|---|---|
| 1 | *Hypsipetes leucocephalus* |
| 2 | *Acritillas indica* |
| 3 | *Alophoixus ruficrissus* |
| 4 | *Setornis criniger* |
| 5 | *Pycnonotus leucogenys* |
| 6 | *Rubigula squamata* |
| 7 | *Phyllastrephus poliocephalus* |
| 8 | *Arizelocichla striifacies* |
| 9 | *Neolestes torquatus* |
